# Quantifying adaptive evolution and the effects of natural selection across the Norway spruce genome

**DOI:** 10.1101/2020.06.25.170902

**Authors:** Xi Wang, Pär K Ingvarsson

**Affiliations:** Umeå Plant Science Centre, Department of Ecology and Environmental Science, Umeå University, SE901 87 Umeå, Sweden; Linnean Centre for Plant Biology, Department of Plant Biology, Swedish University of Agricultural Sciences, SE750 07 Uppsala, Sweden

**Keywords:** *Picea abies*, whole-genome re-sequencing, negative selection, positive selection, balancing selection

## Abstract

Detecting natural selection is one of the major goals of evolutionary genomics. Here, we sequence whole genomes of 34 *Picea abies* individuals and quantify the amount of selection across the genome. Using an estimate of the distribution of fitness effects, we show that negative selection is very limited in coding regions, while positive selection is rare in coding regions but very strong in non-coding regions, suggesting the great importance of regulatory changes in evolution of Norway spruce. Additionally, we found a positive correlation between adaptive rate with recombination rate and a negative correlation between adaptive rate and gene density, suggesting a widespread influence from Hill-Robertson interference to efficiency of protein adaptation in *P. abies*. Finally, the distinct population statistics between genomic regions under either positive or balancing selection with that under neutral regions indicated impact from selection to genomic architecture of Norway spruce. Further gene ontology enrichment analysis for genes located in regions identified as undergoing either positive or long-term balancing selection also highlighted specific molecular functions and biological processes in that appear to be targets of selection in Norway spruce.

## Introduction

Natural selection leaves detectable signatures in the genome of a species and characterizing and quantifying such signatures at the molecular level is one of the major goals of evolutionary genomics. Detecting genes, or genomic regions, that have been targeted by natural selection is significant not only because they illustrate the action of evolutionary processes and shed light on species histories, but also because they could represent biologically meaningful variation that may provide important functional information (Nielsen 2005; Vitti et al. 2013).

The nearly neutral theory (Ohta 1973) was proposed as an extension to the neutral theory model (Kimura 1983) to overcome some of the shortcomings of the neutral model which inadequately explained emerging molecular data, in particular the constancy of the molecular clock (Chen et al. 2020). Compared with the neutral theory, where mutations are assumed to be either neutral or strongly deleterious, the nearly neutral theory also considers a class of mutations that are weakly selected and effectively neutral and the fraction of mutations affected by selection hence depends on the effective population size (Ohta 1973; Ohta 1992; Ohta and Gillespie 1996). This weak selection model was best described by Kreitman (1996) as “the slightly deleterious model” and primarily considers slightly deleterious mutations. Both of the strict neutral theory and the nearly neutral theory assume that “only a minute fraction of DNA changes in evolution are adaptive in nature” (Kimura 1983; Ohta 1973) and thus suggest that mutations influenced by positive selection are too rare to have any “statistically” significant effect on the rate of evolution in most organisms (Ingvarsson 2010). Alternative models that allocate a greater role to positive selection have been proposed and these suggest that the rate of evolution should be determined by both the beneficial mutation rate and how quickly these mutations can spread and ultimately fix in a species (Gillespie 1991; Gillespie 2000; Gillespie 2001). Under such models, clarifying and quantifying the relative contribution of neutral, beneficial, and deleterious mutations to rates of evolution across the genome is one of the outstanding problems in evolutionary genetics.

Based on “the slightly deleterious model”, Keightley and Eyre-Walker (2007) developed a site frequency spectra (SFS) based maximum-likelihood approach that combine within-species nucleotide polymorphism data and parameters of a demographic model that allows a population size change at some time in the past, to estimate the distribution of fitness effects (DFE) of newly arisen mutations. Based on this method, the amount of negative selection acting on different categories of sites can be estimated and quantified by comparing two sets, one set assumed to be evolving neutrally and subject to the action of mutation and genetic drift and another set of sites assumed to be evolving under the influence of natural selection (Keightley and Eyre-Walker 2007). The method was subsequently extended to also estimate the efficiency of adaptive molecular evolution by inferring the rate and fitness effects of advantageous mutations by accounting for the contribution of slightly deleterious mutations to polymorphism and divergence (Eyre-Walker and Keightley 2009). This method thus makes it possible to quantify the amount of positive selection acting in species and thus to test models that allocate a greater role to positive selection (Gillespie 1991; Gillespie 2000; Gillespie 2001) while still regarding the nearly neutral model as the *de facto* null model.

Current evidence suggests that both positive and negative selection are common in coding and some noncoding regions in several model systems, e.g. *Drosophila* (Andolfatto 2005; Haddrill et al. 2008; Fraïsse et al. 2019), humans (Torgerson et al. 2009; Lindblad-Toh et al. 2011; Arbiza et al. 2013), and mice (Halligan Det al. 2010; Kousathanas et al. 2011). It has been suggested that the majority of adaptive evolution may occur in noncoding, regulatory regions, because new mutations that occur in these regions may have fewer deleterious pleiotropic effects (Carroll 2005; Wray 2007). Halligan et al. (2013) showed that there have been many more adaptive substitutions in noncoding DNA than in coding regions in house mice, although adaptive substitutions in coding regions may experience stronger positive selection. However, our understanding of the action of natural selection remains relatively limited in plants, particularly in noncoding regions. Recently, studies in *Capsella grandiflora* found widespread positive and negative selection in both coding and regulatory regions, but also suggested that both positive and negative selection on plant noncoding sequences are considerably rarer than in animal genomes (Williamson et al. 2014).

Evidence has accumulated over the last decade that show that rates of adaptive protein evolution are high in some species. For instance, 45% of all amino acid substitutions are thought to have been fixed by positive selection in Drosophila (*D. simulans* and *D. yakuba*, Smith and Eyre-Walker 2002), more than 50% in enteric bacteria (*E. coli* and *S. enterica*, Charlesworth and Eyre-Walker 2006), 57% in wild mice (*Mus musculus castaneu*s, Halligan et al. 2010), 40% in *C. grandiflora* (Slotte et al. 2010), 30% in *Populus tremula* (Ingvarsson 2009), and between 10-20% of substitutions differentiating humans and chimpanzees (Gojobori et al. 2007; Boyko et al. 2008). Estimating rates of adaptive evolution in protein coding sequences and the findings that rates of adaptive evolution differs across species opens up possibilities to investigate the factors affecting the efficiency of natural selection. Hill-Robertson interference (HRi) is expected to reduce the overall efficiency of natural selection when there is linkage between sites under selection (Hill and Robertson 1966; Felsenstein 1974; Comeron et al. 2008; Castellano et al. 2016). Specifically, when a newly arisen advantageous mutation is linked other beneficial mutations, the probability of fixation will be reduced because of competition among the different adaptive mutations. Similarly, when an advantageous mutation arise in linkage disequilibrium with deleterious mutations, its fixation probability will also decrease if it cannot recombine away from the deleterious background (Comeron et al. 2008; Castellano et al. 2016). The magnitude of this linkage effect depends on recombination rates and the strength of selection. We therefore expect a positive correlation between recombination rate and the rate of adaptive evolution, as the influence of linkage is expected to be stronger, and will hence result in stronger HRi, in regions of low compared to high recombination. Similarly, genes embedded in gene rich regions should show stronger HRi due to stronger linkage effect than genes located in gene poor regions, because the densities of selected sites are thought to be higher in gene-rich regions. The net result is an expected negative correlation between gene density and rates of adaptive evolution. Finally, genes with high mutation rates are predicted to adapt faster than those with low mutation rates, regardless of whether most adaptation comes from newly arising mutations or from standing genetic variation (Castellano et al. 2016). This prediction is expected in species where the overall mutation rate is low since natural selection remains effective and increasing the mutation rate can give rise to an increase in the rate of adaptation substitutions. However, when the mutation rate in a species is high to begin with, increasing the mutation rate may have a detrimental effect because of the overwhelming majority of novel mutations are expected to be deleterious (Gerrish et al. 2013).

The factors that maintain genetic and phenotypic variation within natural populations have long been an important topic in evolutionary biology (Delph and Kelly 2014). Positive selection raise the frequency of adaptive mutations over time in a population while negative selection decreases the frequency of alleles that impair fitness and both processes acts to reduce genetic diversity (Dutheil 2020). In contrast, balancing selection maintains multiple advantageous polymorphisms in populations, leading to an increased genetic diversity in regions surrounding a balanced polymorphism (De Filippo et al. 2016). Evidence of balancing selection have been investigated in many species, e.g. Koenig et al. (2019) pointed to long-term balancing selection as an important factor shaping the genetics of immune systems in plants and as the predominant driver of genomic variability after a population bottleneck, by documenting genomic variability after two parallel species-wide bottlenecks in the genus *Capsella*. Wang et al. (2019) provided evidence that, apart from background selection, both recent positive selection and long-term balancing selection have also been crucial components in shaping patterns of genome-wide variation during the speciation process among the three aspen species (*P. tremula, P. davidiana* and *P. tremuloides*) by inferring the genealogical relationships and estimating the extent of ancient introgression across the genome.

Conifers are the most widely distributed group of gymnosperms and are estimated to cover ∼39% of the world’s forests (De La Torre et al. 2014). Confers genomes are large (typically 20-40 Gb) and highly repetitive but nevertheless show a largely conserved synteny even over long evolutionary timescales (Pavy et al. 2012, Nystedt et al 2013). The large size of most conifer genomes have made them inaccessible to genome-wide studies, but the recent publication of a draft reference genome for Norway spruce (*Picea abies*, Nystedt et al 2013) has opened up possibilities for whole-genome resequencing in this species and thus also to assess how patterns of natural selection vary across the genome (Wang et al. 2020). In this paper, we use available genome resources for Norway spruce together with whole-genome resequencing data generated from a sample of trees spanning the distribution range of *P. abies* to quantify the strength of both positive and negative selection acting on coding and non-coding regions. In addition, we use genome scans across the Norway spruce genome to identify genomic regions that have been targets by either recent positive and long-term balancing selection to assess how natural selection affects patterns of variation and to also understand the functions of gene located in those regions.

## Methods

### Sampling, sequencing, and variant calling

We used whole genome re-sequencing data from 34 individuals of Norway spruce (*P. abies*) previously described in Wang et al. (2020). The samples were collected to span the natural distribution of the species, extending from Finland and Russia in the east to Sweden and Norway in the west and Belarus, Poland, Romania in the south (supplementary Table S1; supplementary Figure S1). All samples were collected from newly emerged needles or dormant buds and were stored at -80 °C until DNA extraction using a Qiagen plant mini kit following manufacturer’s instructions. All sequencing was performed at the National Genomics Initiative platform at the SciLifeLab facilities in Stockholm, Sweden, using paired-end libraries with an insert size of 500 bp.

The bioinformatics pipeline used to handle all sequencing data has previously been described in detail in Bernhardsson et al. (2020) and Wang et al (2020). Briefly, all raw sequence reads were first mapped to the complete *P. abies* reference genome v.1.0 (Nystedt et al. 2013) using BWA-MEM v0.7.15 (http://bio-bwa.sourceforge.net/bwa.shtml, Li 2013) with default settings. To reduce the computational complexity of the subsequent SNP calling (due to the large genome size, high repetitive content and fragmented genome assembly), we reduced the data by only considering genomic scaffolds greater than 1kb in size genome and then subdivided the BAM files containing the mapped reads into 20 smaller data subsets to enable us to curate the data in parallel and to enable existing software tools (e.g GATK) to handle the entire data set. In order to eliminate artifacts introduced due to DNA amplification by PCR, which could potentially lead to excessively high read depth in some regions, PCR duplicates were marked in all data subsets using MarkDuplicates in Picard v2.0.1 (http://broadinstitute.github.io/picard/). Local realignment was further performed to minimize mismatching bases occurring in regions with insertions and/or deletions (indels) during the mapping step by first flagging suspected intervals using RealignerTargetCreator, followed by realignment of those intervals using IndelRealigner, both implemented in GATK v3.7 (DePristo et al. 2011). Finally, we performed variant calling using GATK HaplotypeCaller to generate intermediate genomic VCFs (gVCFs) and then carried out joint calling on all gVCF files for the 34 samples using the GenotypeGVCFs module in GATK. Information including original sampling location, platform used, and estimated coverage from raw sequencing reads and of BAM files after mapping are given for all individuals in supplementary Table S1. In addition, we downloaded raw sequences from one individual of White spruce (*P. glauca*) to use as an outgroup from the NCBI Sequence Read Archive (https://www.ncbi.nlm.nih.gov/sra?LinkName=bioproject_sra_all&from_uid=242552) and performed all steps of variant calling described above except for the joint calling step.

### Filtering to maintain high quality SNPs

In order to retain high quality SNPs, we performed the following filtering steps for all called variants to reduce the number of false-positive SNPs. We only included biallelic SNPs positioned >5 bp away from an indel and where the SNP quality parameters fulfilled GATK recommendations for hard filtering (https://gatkforums.broadinstitute.org/gatk/discussion/2806/howto-apply-hard-filters-to-a-call-set). We also recoded genotype calls with a depth outside the range 6-30 and a GQ < 15 to missing data and filtered each SNP for being variable with an overall average depth in the range of 8-20 and a “maximum missing” value of 0.8 (max 20% missing data). Finally, as SNPs called in collapsed regions in the assembly, likely containing nonunique regions in the genome, should show excess heterozygosities as they are based on reads that are derived from different genomic regions, we removed all SNPs that displayed a P-value for excess of heterozygosity <0.05. SNPs that passed all the different hard filtering criteria were used in the downstream analyses. For more detailed information on the genotype hard filtering criteria used, please refer to Bernhardsson et al. (2020) and Wang et al. (2020).

### Population structure and estimation of effective population size

We used fourfold synonymous SNPs to perform population structure analyses with PCAngsd by using a method for inferring population structure through PCA using an iterative heuristic approach of estimating individual allele frequencies (Meisner and Albrechtsen 2018). NGS provide large amounts of sequencing data but are also associated with statistical uncertainty, especially for low-coverage data (Meisner and Albrechtsen 2018). A model-based approach can account for this uncertainty by working directly on genotype likelihoods of the unobserved genotypes, improving accuracy in samples with low and variable sequencing depth for datasets.

We calculated the effective population size (N_e_) for each population based on levels of polymorphism quantified by Watterson’s estimator of 4N_e_μ (θ_W_) (Watterson 1975) and nucleotide diversity, π (Tajima 1983) for synonymous sites (Gossmann et al. 2010). The effective population size using the level of synonymous site diversity and dividing this by an estimate of the mutation rate per generation (N_e_= π_synonymous_/4μ), where synonymous site diversity for each population refer to Wang et al. (2020), and a mutation rate of 1.1×10^−9^ per site per year and a generation time of 25 years was used for *P. abies* (Nystedt et al 2013; Chen et al. 2019).

### Functional sites and Genomic Features Estimates: Nucleotide Diversity, Tajima’s D, Divergence, Recombination Rate, Gene Density, GC Density, and Repeat Density

BED files for different genomic contexts (4-fold synonymous sites, 0-fold nonsynonymous sites, introns, and intergenic sites) were generated from the genome annotation for *P. abies* v1.0 (available from ftp://plantgenie.org/Data/ConGenIE/Picea_abies/v1.0/GFF3/) using a custom-made python script (https://github.com/parkingvarsson/Degeneracy/). Separate VCF files were generated for the different site categories from the original VCF files based on the genomic BED files using vcftools.

We used ANGSD v0.921 (Korneliussen et al. 2014) to estimate pairwise nucleotide diversity and Tajima’s D by calculating the site allele frequency likelihood based on normalized phred-scaled likelihoods of the possible genotypes (PL tag in the VCF file). Divergence was calculated between *P*.*abies* and the outgroup species *P. glauga* at 4-fold, 0-fold, intronic, and intergenic sites by measuring the number of fixed differences per scaffold. The population scaled recombination rate (4N_e_r) was estimated per scaffold for each population using a Bayesian reversible-jump Markov Chain Monte Carlo scheme under the crossing-over model as implemented in LDhat v2.2 (McVean et al. 2004). We performed 1,000,000 Markov Chain Monte Carlo iterations with sampling every 2,000 iterations and set up a block penalty parameter of 5 using a data set consisting of only scaffolds longer than 5 kb because shorter scaffolds generally did not produce stable estimates. The first 100,000 iterations of the reversible-jump Markov Chain Monte Carlo scheme were discarded as a burn-in. We measured gene density per scaffold as the ratio of sites falling within a gene model on the scaffold to the overall length of the scaffold. The same method was used to estimate repeat density using information on repeat content per scaffold (ftp://plantgenie.org/Data/ConGenIE/Picea_abies/v1.0/GFF3/Repeats/). GC density was calculated at the scaffold level as the fraction of bases where the reference sequence (*P. abies* genome v1.0) was a G or a C using BEDtools.

### Estimating the distribution of deleterious fitness effects and the fraction of adaptive substitutions for both genic and non-genic regions

The distribution of fitness effects of new mutations (DFE) specifies the probability of a new mutation having a given fitness effect (Keightley and Eyre-Walker 2007). The software DFE-alpha (Keightley and Eyre-Walker 2007) was employed to estimate the fraction of sites under negative selection with different effective strengths by incorporating the expected allele frequency distribution generated by transition matrix methods and simultaneously a demographic model that includes a step population size change. This method was based on the “slightly deleterious model” which assume that the fitness effects of new mutations at putatively neutral sites are zero and that mutations are unconditionally deleterious at selected sites as advantageous mutations are assumed to be too rare to contribute to polymorphism (Keightley and Eyre-Walker 2007). In order to make the results are more robust, we generated the folded SFS for each category of selected sites (0-fold nonsynonymous sites, intronic, and intergenic sites) and a class of putatively neutral reference sites (4-fold synonymous sites) from SNP data using ANGSD and report the proportion of mutations falling into different effective strengths of selection (N_e_s, where N_e_ is the effective population size and s is the selection coefficient) range: 0-1, 1-10, 10-100, and >100, which corresponding to effectively neutral, mildly deleterious, deleterious, and strongly deleterious, respectively.

Based on the estimated distribution of fitness effects of new deleterious mutations from the polymorphism data, DFE-alpha further predict the numbers of substitutions originating from neutral and slightly deleterious mutations between two species by using divergence data (Eyre-Walker and Keightley 2009). If the observed number of substitutions is greater than this expectation, the proportion of adaptive substitutions (α) and the rate of adaptive substitutions expressed relative to the neutral substitution rate (ω) is estimated from the difference between observed and expected substitutions (Eyre-Walker and Keightley 2009). We used *P. glauga* as outgroup to infer parameters under positive selection at 0-fold, intronic, and intergenic sites of Norway spruce genome. Jukes-Cantor multiple hits correction was applied to the divergence estimates (Jukes and Cantor 1969). For all parameters of N_e_s, α, and ω, we generated 100 bootstrap replicates by resampling randomly across all scaffolds for each site class using R (R Develpment Core Team 2017). Confidence intervals of 95% for each parameter was represented by excluding the top and bottom 2.5% of bootstrap replicates.

### Gene Bins and the factors affecting amino acid adaptation

In order to understand the factors influencing the efficiency of natural selection, we further assessed correlations between the rate of adaptive evolution within genic regions, introns and intergenic regions. To avoid the noisy and large sampling variances arising from estimates of single genes due to limited numbers of segregating or divergent sites for some site classes (Stoletzki and Eyre-Walker 2011, Castellano et al. 2016; Moutinho et al.2019), we grouped scaffolds with genes into bins according to their estimated rate of recombination, gene density and mutation rates. The rank of values for all these bins can be referred in the supplementary Table S3-S4. The rate of adaptive evolution can be estimates in two different ways, either as the rate of adaptive evolution relative to the neutral substitution rate (ω) or through the rate of adaptive amino acid substitutions (Ka+). However, ω is not useful for studying the effects of the mutation on the rate of adaptive evolution because it is normalized by the mutation rate and we therefore focused on assessing correlations between Ka+ and the rate of recombination, gene density and mutation rate, respectively. In this study, assuming that 4-fold synonymous sites are free of the effects of selection and thus serves as our baseline, we first detected the proportion of adaptive substitutions (α) (0-fold non-synonymous sites) in each bin by running DFE-alpha (Eyre-Walker and Keightley 2009). From these estimates, Ka+ can be further estimated using the expression: Ka+ = αKa, where Ka represents the number of non-synonymous substitutions per non-synonymous site. As a comparison, we also calculated the ratio of non-synonymous to synonymous substitutions (K_a_/K_s_) for all bins separately and estimate the correlation between the K_a_/K_s_ ratio and the genomic features of interest.

When analyzing the role of the mutation rate for adaptation, we estimated correlations between the rate of adaptive substitutions at 0-fold non-synonymous sites (Ka+) and 4-fold synonymous divergence (K_4_) from the outgroup *P*.*glauga*. To overcome the limitation that the estimation of Ka+ depends on K_4_, we split K_4_ into three independent variables using sampling without replacement, following ideas described in several recent papers (Smith and Eyre-Walker 2002; Piganeau and Eyre-Walker 2009; Stoletzki and Eyre-Walker 2011; Gossmann et al. 2012; Castellano et al. 2016). The first K_4_ variable (K_4,1_) was used to rank scaffolds with genes, while the second variable (K_4,2_) was used to estimate the rate of 0-fold adaptive substitutions (Ka+) and the last variable (K_4,3_) was regarded as an estimate of the mutation rate.

All statistical analyses were performed using the R statistical package (R Core Team 2017). Linear regressions were carried out with the R function “lm” and nonlinear regression were run using the R function “nls”. To compare the linear and non-linear model fits, we calculated Akaike’s AIC using the R functions “AIC” and “BIC”. Pairwise correlations between the variables of interest were calculated using Spearman’s rank correlations using the basic R function “cor.test”.

### Genome-wide scan for regions under positive selection and balancing selection

Selective sweeps leave distinct signatures in the genome or an organism (Stephan 2019). To identify the loci and/or regions that have undergone recent positive selection, we scanned the whole genome of Norway spruce, using RAiSD (Raised Accuracy in Sweep Detection), which use multiple signatures of a selective sweep via the enumeration of SNP vectors (Alachiotis and Pavlidis 2018). This program introduced the *μ* statistic, a composite evaluation test that scores genomic regions by quantifying changes in the SFS, the levels of LD, and the amount of genetic diversity along a chromosome, achieving high sensitivity and accuracy while reducing the computational complexity, which allows for faster processing with limited memory requirements (Alachiotis and Pavlidis 2018). We ran RAiSD using default settings on a subset of the complete data, consisting of individuals from the Sweden-Norway population (25 individuals), to remove possible influences due to population structure.

In order to detect regions under long-term balancing selection, we scanned the whole genome of Norway spruce (again using the Sweden-Norway population to limit the effects of population structure) using the *β* (beta) score summary statistic which detects clusters of alleles at similar frequencies (Siewert and Voight 2017). This statistic was proposed based on simulations showing that new mutations which arise in close proximity to a site targeted by balancing selection accumulate at frequencies nearly identical to that of the balanced polymorphism (Siewert and Voight 2017). Compared to existing summary statistics, the *β* score has improved power to detect balancing selection, is reasonably powered under non-equilibrium demographic models and across a range of recombination and mutation rates, and is computationally efficient and applicable to species that lack appropriate outgroup sequences (Siewert and Voight, 2017). We first converted our VCF file to a betascan input file by running two scripts, “vcfm2acf” and “acf2betascan”, from the glactools package (https://github.com/grenaud/glactools). We calculated the folded *β* statistic in 1 kbp windows since the signals of long-term balancing selection are usually localized to very narrow genomic regions (Gao et al, 2015; Wang et al. 2020). To prevent false positives, we filtered out SNPs with a folded frequency lower than 20%, and defined significant SNPs as those SNPs with extreme *β* scores in the top 0.1% of sites from the genome-wide *β* score distribution.

To assess the effects of positive and balancing selection on patterns of genome-wide variation, we compared outlier windows identified in two methods, representing regions under positive or balancing selection, with the remaining genomic regions using a variety of population genetic summary statistics, including genetic diversity per site (π), Tajima’s D, the population scaled recombination rate, gene density, GC density and repeat density for Sweden-Norway population (25 individuals).

### GO Enrichment

To determine whether any functional category were overrepresented among genes in the regions that we identified as being under positive or balancing selection, we performed functional enrichment analysis of GO categories using Fisher’s exact test (http://congenie.org/enrichment). P-values for Fisher’s exact test were further corrected for multiple testing using the Benjamini-Hochberg FDR method (Benjamini and Hochberg 1995). GO terms with an FDR corrected P-value <0.1 were considered to be significantly enriched.

## Results and Discussion

Whole-genome resequencing data were generated for 34 individuals sampled to span the natural distribution of *P. abies* using Illumina HiSeq 2000 or HiSeq X with a mean sequence coverage of 18.1x per individual (supplementary table S1). All raw sequence reads were mapped to the complete *P. abies* reference genome v.1.0, which contains ∼10 million scaffolds covering 12.6Gb out of the estimated genome size of ∼20Gb (Nystedt et al. 2013). To reduce the computational complexity of SNP calling, BAM files were subsetted to only include scaffolds longer than 1kb and then subdivided into 20 genomic subsets with ∼100,000 scaffolds in each. After PCR duplication removal, local realignment, variant calling and several steps for hard filtering, 293.9 million high quality SNPs remained for all downstream analyses.

### Population structure

In order to account for the effects arising from population structure, we performed a PCA analysis though an iterative heuristic approach of estimating individual allele frequencies by PCAngsd (Meisner and Albrechtsen 2018). Based on 4-fold synonymous sites using data for 33 individuals (Pab034 was removed because it was found to be highly related with Pab033, see Wang et al. 2020 for more details), the PCA result clearly reflects geographic structure within *P. abies* (supplementary Figure S2): samples from Belarus, Poland, and Romania grouped into one cluster, most Finish samples clustered into one group, most individuals from Norway and northern Sweden clustered into one group. Two individuals, one from southern Finland (Pab015) and one from southern Sweden (Pab002), could not be grouped within any of the populations as they fell between the three main groups in the PCA plot, indicating that they represent materials that have recently been introduced in to Sweden and Finland from elsewhere in Europe (Chen et al. 2019; Wang et al. 2020). We therefore removed these individuals (Pab015 and Pab002) from further analyses and regarded remaining samples as being derived from three populations corresponding to the groupings seen in the PCA, hereafter referred to as “Central-Europe,” “Finland,” and “Sweden-Norway” (see also Wang et al 2020). Earlier results have shown that *P. abies* can be subdivided into three main domains, including the Alpine domain, the Carpathian domain, and the Fennoscandian (Baltico-Nordic) domain (Heuertz et al. 2006; Chen et al. 2019). Our population structure analyses suggest that our Central-Europe population is likely composed of individuals derived from the Carpathian and/or Alpine domains, and there is also sub-structure within the Fennoscandian (Baltico-Nordic) domain that could represent either population structure within the Fennoscandian domain or the effects of ongoing hybridization with *P. obovata* that is more apparent in the eastern (Finland) part of the distribution range (Tsuda et al 2016). Both the results from the PCA in this study and previous research thus suggest a relatively strong population structure of Norway spruce across its range.

### Genome-wide measures of purifying selection

Purifying selection in a genomic region is usually considered to be a clear sign that the regions has some functional importance and therefore show evolutionary conservation. In order to quantify the amount and strength of purifying selection acting across the genome of Norway spruce, we used the methods of Keightley and Eyre-Walker (2007) to infer the percentage of deleterious mutations falling into different categories based on the effective strength of negative selection (in terms of N_e_s). We assessed this for a number of different categories of sites, including 0-fold non-synonymous, intronic and intergenic sites using 4-fold synonymous sites as a putatively neutral baseline (Figure 1A; Table 1). For 0-fold non-synonymous sites, most of the mutations fell in the range representing weakly deleterious mutations that behave as effectively neutral (0<N_e_s<1), making up 68.5% (95% CI: 61.9%-77.7%) of all sites. Similarly, 13.9% (95% CI: 3.80%-22.1%) of amino acid mutations are considered strongly deleterious (N_e_s>100), suggesting that they are under strong purifying selection. The remaining non-synonymous mutations are under moderate level of negative selection, of which 8.4% (95% CI: 7.5%-9.3%) are classified as mildly deleterious (1<N_e_s<10) and 9.3% (95% CI: 8.5%-9.6%) are deleterious mutations (10<N_e_s<100). As a comparison, for both intronic and intergenic sites, the vast majority mutations, approaching 100%, are considered nearly neutral with N_e_s falling in the range between 0-1.

**Table 1.** Estimates of the distribution of fitness effects of new mutations at zerofold nonsynonymous sites, intronic sites, and intergenic sites falling in different N_e_s ranges, and proportion of divergence driven to fixation by positive selection (α) and the rate of adaptive substitution relative to neutral divergence (ω) in *P. abies*

| *P. abies* | Negative Selection |  |  |  | Positive Selection |  |
| --- | --- | --- | --- | --- | --- | --- |
| Category | $N_e$s<br>(0–1) | $N_e$s<br>(1–10) | $N_e$s<br>(10–100) | $N_e$s<br>(>100) | *P. glauca* (outgroup) | |
| | | | | | $\alpha$ | $\omega$ |
| Zero-fold | 0.685<br>(0.619-0.777) | 0.084<br>(0.075-0.093) | 0.093<br>(0.085-0.096) | 0.139<br>(0.038-0.221) | 0.099<br>(-0.073-0.221) | 0.073<br>(-0.045-0.175) |
| Intronic | 1.000<br>(1.000-1.000) | 0.000<br>(0.000-0.000) | 0.000<br>(0.000-0.000) | 0.000<br>(0.000-0.000) | 0.417<br>(0.303-0.501) | 0.716<br>(0.435-1.003) |
| Intergenic | 1.000<br>(1.000-1.000) | 0.000<br>(0.000-0.000) | 0.000<br>(0.000-0.000) | 0.000<br>(0.000-0.000) | 0.139<br>(0.005-0.232) | 0.161<br>(0.005-0.302) |

Ninety-five percent bootstrap confidence intervals are shown in parentheses.

**Figure 1.**
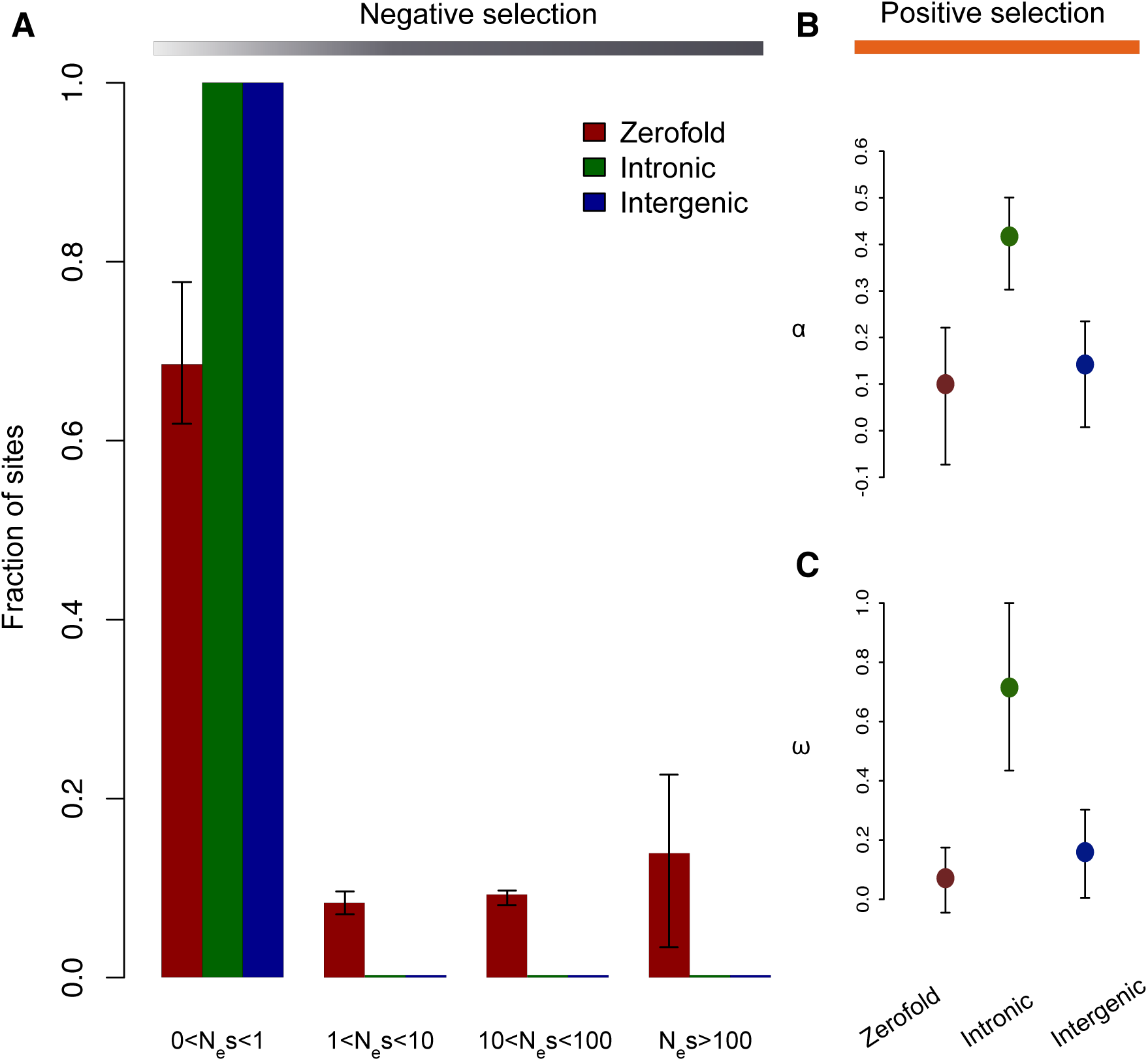
Estimates of negative and positive selection on coding and noncoding sites in *P. abies*. (A) The proportion of sites found in each bin of purifying selection strength, separated by site type. (B) The proportion of divergent sites fixed by positive selection, and (C) the rate of adaptive substitution relative to neutral divergence. Error bars represent 95% bootstrap confidence intervals.

By analyzing more than 2,400 loci with an average length of ∼280 nucleotides from 11 plant species, Gossmann et al. (2010) estimated the distribution of fitness effects of new mutations (DFE) in protein-coding sequences and found that in all species, the largest proportion of mutations are strongly deleterious with N_e_s>100 and for most species less than 25% of amino acid-changing mutations behave as effectively neutral (0 < N_e_s < 1). However, three species (*Boechera stricta, Populus. balsamifera*, and *Schiedea globosa*) showed an excess of neutral mutations (>25%) and a decrease of strongly deleterious mutations (<55%), suggesting a clear impact of the species-wide effective population size (N_e_) on the DFE so that species with small N_e_ tend to have a relatively large proportion of mutations that are effectively neutral. Gossmann et al. (2010) additionally estimated the DFE from both wild and domesticated populations of rice and found that domesticated varieties of *Oryza. japonica* showed a higher proportion of effectively neutral mutations than the wild species *O. rufipogon*, which may reflect a lower effective population size associated with domesticated varieties.

Wang et al. (2020) inferred the demographic history of Norway spruce and found that it is characterized by several reoccurring bottlenecks corresponding to drastic climate fluctuations during the Quaternary with concomitant decreases in effective population size, despite the fact that the species has a widespread current geographic distribution. Historical population size fluctuations and reoccurring bottlenecks have sharply reduced the effective population size in Norway spruce and is one of the likely reasons for the large proportion of new amino acid mutations that we classify as effectively neutral. Human-mediated selection and the use of limited seed sources for reforestation could also have contributed to a reduction in the effective population size. Chen et al. (2019) showed that a large proportion of the 1,499 individuals stemming from the Norway spruce breeding program in southern Sweden corresponds to recent introductions from mainland Europe. This fact suggests that humans have affected the demography of Norway spruce through breeding, although the extent of such processes are unclear and are worthy of further study. Moreover, we only estimated the DFE for deleterious mutations, rather than of the DFE of both deleterious and adaptive mutations (Keightley and Eyre-Walker 2007). Putative mutations under positive selection will be added to the category of effectively neutral mutations, leading to a higher proportion of mutations falling to range representing effectively neutral (0<N_e_s<1), although the biases due to these processes vary with N_e_ (Chen et al. 2017). Finally, the presence of strong and extensive codon usage bias might violate the presumed neutrality of synonymous sites.Several analyses of codon usage bias from population genetic data does suggest the action of selection on synonymous sites in plant species (Ingvarsson 2010; Qiu et al. 2011; Lawrie et al. 2013; Machado et al. 2020). De La Torre et al. (2015) found high levels of codon bias, measured as F_op_, for in *P. abies* and *P. glauca* when analyzing genome-wide levels of gene expression (>50,000 expressed genes) data. Thus, selection on at least a fraction of the synonymous sites in Norway spruce due to strong codon usage bias cannot be ruled out and may therefore contribute to a downwardly biased estimate of the proportion of strongly deleterious amino acid mutations.

Compared to the variable proportions of different classes of purifying selection observed at non-synonymous sites, negative selection appears to be largely absent in both introns and intergenic region. This is similar to what Williamson et al. (2014) found using data from whole genome sequences for 13 *Capsella grandiflora* individuals. Their data showed that the proportion of intergenic sites that are nearly neutral approached 100% and that approximately 70% of intronic sites were behaving as effectively neutral. Further, after bootstrapping the latter estimate was deemed not significantly different from 100% of intronic sites being effectively neutral. However, considering only conserved noncoding sequences (CNSs), Williamson et al. (2014) found that at least 5% of sites in intergenic regions are under strong purifying selection. Although we classified all non-coding sites as falling in the nearly neutral category in Norway spruce, the DFE approach might not sensitive enough to detect negative selection on a very small subset of sites. Similar patterns were also found by Lin et al. (2019) which showed that negative selection was rare or absent in non-coding regions for two aspen species, with 72.7% of introns sites in *Populus tremula*, 74.8% of intron sites in *P. tremuloides* and 100% in intergenic region for both species falling in the effectively neutral category (0<N_e_s<1). However, our results for Norway spruce are in stark contrast with results from Drosophila and from humans, where a relatively large fraction of sites in non-coding regions are under selection. For example, in Drosophila, only 30%–70% of intronic and intergenic regions are classifies as nearly neutral (Andolfatto 2005; Sella et al. 2009; Hough et al. 2013; Williamson et al. 2014). Finally, the large proportion of repetitive DNA located in non-genic regions in Norway spruce (Nystedt et al. 2013) may help explain why the overwhelming majority of non-coding sites are falling in the nearly neutral class when we estimate negative selection, despite the fact that many of these sites appear to evolve under positive selection through, for instance, regulatory changes (see below).

### Genome-wide measures of positive selection

The extent to which positive selection contributes to molecular evolution has been a long-standing question in evolutionary genetics (Booker et al. 2017). By taking into account the effect of slightly deleterious mutations, we employed an extension of the McDonald-Kreitman test (Eyre-Walker and Keightley 2009), to estimate the proportion of adaptive substitutions (α) and the rate of adaptive substitutions expressed relative to the neutral substitution rate (ω). For these analyses we used *P. glauca* as outgroup and focused on estimating rates of adaptive evolution at 0-fold sites and sites in intronic and intergenic regions across the Norway spruce genome. The results suggest that non-coding regions show relatively high proportions adaptive substitutions, with intron sites having the highest proportion of adaptive substitutions (α=0.417) and the highest estimate of the rate of adaptive substitutions (ω=0.716). Intergenic sites have substantially lower proportions of adaptive substitutions (α=0.139) as well as rate of adaptive substitutions (ω=0.161, Figure1B; Figure1C; Table1). Bootstrap analysis for both the proportion of adaptive substitutions and adaptive rate (Figure1B; Figure1C; Table 1) showed that these estimates are significantly greater than 0 for both intronic and intergenic sites, suggesting the action of widespread positive selection in non-coding regions in Norway spruce. As a comparison, we found the lowest proportion of adaptive mutations (α=0.099) and lowest rate of adaptive evolution (ω=0.073) at 0-fold non-synonymous sites (Figure 1B; Figure 1C; Table1).

It has been suggested that the majority of adaptive evolution may occur in noncoding regulatory regions, where new mutations may have fewer deleterious pleiotropic effects (Carroll 2005; Wray 2007). Andolfatto (2005) assessed the mode of selection acting on non-coding DNA in *Drosophila*, using polymorphism data for 35 coding fragments (average length 667 base pairs) and 153 noncoding fragments (average length 426 bp) scattered across the X chromosome of *D. melanogaster*. By comparisons with the closely related sibling species, *D. simulans*, Andolfatto (2005) showed that a substantial fraction of the nucleotide divergence in non-coding regions had been driven to fixation by positive selection (about 20% for most intronic and intergenic DNA, and 60% for UTRs), suggesting that adaptive changes in non-coding DNA might have been considerably more common in the evolution of *D. melanogaster* than earlier thought. A significant role for adaptive evolution in non-coding regions has also been reported in other species. For instance, at least 1MB of functional sequence has accumulated many beneficial nucleotide replacements within the non-coding fraction of the human genome (Ponting et al. 2006) and approximately 80% of all adaptive substitutions in the mouse genome are located in noncoding regions (Halligan et al. 2013). Our results show that a greater proportion of adaptive substitutions occur in non-coding regions compared to coding regions in the Norway spruce genome, supporting previous suggestions about the importance of regulatory changes in evolution (King and Wilson 1975; Carroll et al. 2001). As a comparison, Williamson et al. (2014) showed that there is little evidence for a difference in the strength of positive selection on substitutions in coding regions compared to noncoding regions in the flowering plant *C. grandiflora*. This observation is consistent with earlier suggestions that, unlike in animals, plant genomes may contain relatively few noncoding regulatory sequences that are subject to selection, possibly because gene expression can be modified through frequent gene duplication and functional divergence rather than through the evolution of novel regulatory elements (Lockton and Gaut 2005). The very large proportion of adaptive substitutions and the high adaptive rate of evolution we observe in non-coding regions in Norway spruce genome therefore require a novel explanation. Polyploidy is a common mode of speciation and evolution in angiosperms (Leitch and Bennett 1997), however, there is no evidence for recent whole-genome duplications in the gymnosperm lineage (Nystedt et al. 2013). Thus, gene duplications might be rare in the Norway spruce genome possibly suggesting more important roles for novel regulatory elements located in non-coding regions for modifying gene expression. Moreover, conifer genomes contain an abundance of repeat-rich content, mostly in the form of transposable elements (De La Torre et al. 2014). Genomic repeats, and in particular transposable elements, has been a rich source of material for the assembly and tinkering of eukaryotic gene regulatory systems (Feschotte 2008). The very large repetitive fraction (>70%) of Norway spruce genome, and specifically the fraction of Long terminal repeat-retrotransposons (LTR-RTs) comprising the Ty3/Gypsy superfamily and Ty1/Copia superfamily, make contributions to large proportion of putative regulatory elements that ultimately could result in higher rates of adaptive evolution in non-coding regions compared to coding regions.

### Determinants of the efficacy of amino acid adaptive evolution

Compared to the high rates of adaptive evolution we observe in non-coding regions, the fraction of adaptive substitutions fixed by positive selection and the scaled rate of adaptive evolution at 0-fold non-synonymous sites are low and not significantly different from zero as judged by the bootstrap estimates (α 95% CI: -0.073-0.221 and ω 95% CI: - 0.045-0.175, Figure1B; Figure1C; Table 1). This suggests that the proportion and rate of adaptive evolution in coding regions in Norway spruce are limited. Using DNA sequence data from 167 orthologous nuclear gene fragments, Eckert et al. (2013) found little evidence for long-term adaptive non-synonymous evolution in 11 species of soft pines (subgenus *Strobus*), with only one of the α estimates being significantly different from zero. It is reasonable to expect that estimates of α will vary among plants with differing life histories, because life history characteristics of plants are known to affect standing levels of genetic diversity (Eckert et al. 2013). Soft pines are important components of coniferous forests distributed throughout the Northern Hemisphere and they share many life history characteristics with Norway spruce, such as long generation time and a predominantly outcrossing mating system that rely almost exclusively on wind-pollination (Neale and Wheeler 2019), and it is thus nor surprising that we observe similar patterns of adaptive evolution in soft pines and Norway spruce. As a contrast, some angiosperm plants have reported relatively large proportions of amino acid substitutions being fixed by positive selection, for example 40% in *C. grandiflora* (Slotte et al. 2010) and 75% in sunflowers (Strasburg et al. 2009). Even angiosperm species that have similar life history traits as gymnosperms may still show much higher rates of adaptive evolution. Estimates in species from the genus *Populus* that are long-lived outcrossing forest trees show proportions of amino acid substitutions being fixed by positive selection in excess of 30% in *Populus tremula* (Ingvarsson 2010) and 33.8% in *P. tremuloides* (Lin et al. 2019), suggesting life history traits may not play a large role in determining adaptive evolution but rather other factors. An open question that thus remains is why we observe such a large disparity in the rates of adaptive evolution at non-synonymous sites among different plant species and what factors are important for explaining variation in the efficacy of selection.

Hill-Robertson interference (HRi) is expected to reduce the overall efficiency of natural selection when a newly arisen advantageous mutation occurs in linkage disequilibrium with either other beneficial mutations or with deleterious mutations (Hill and Robertson 1966; Felsenstein 1974; Comeron et al. 2008; Castellano et al. 2016). The magnitude of this linkage effect depends on local recombination rate and selection intensities. We therefore first assessed the relationship between the rate of adaptive evolution (Ka+) and recombination rate, expecting to see a positive correlation because the influence of linkage is expected to be stronger, hence resulting in stronger HRi, in regions of low compared to high recombination rates. To estimate the rate of adaptive evolution it is necessary to combine data from several genes because estimates tend to be error prone and sometimes undefined for individual genes (Castellano et al. 2016). We grouped genes into 10 bins based on recombination rates estimated from the Sweden-Norway population, with each bin on average containing 3942 scaffolds and 4634 genes (supplementary table S3). We found a positive correlation between the rate of adaptive evolution and the recombination rate (Spearman’s rank correlation coefficient r =0.236, Figure 2B). This correlation is not significant, likely due to the limited number of bins used and possibly also because estimates population-scaled recombination rates from data on linkage disequilibrium (LD) are unstable in such a fragmented genome assembly as in Norway spruce. We further test whether a curvilinear relationship fits the data better than a linear model, by fitting the function y =a^bx^ to our data and comparing it to the fit of a linear model (supplementary Table S2). Both Akaike’s ‘An Information Criterion’ (AIC) and Schwarz’s Bayesian criterion (BIC) showed that the curvilinear model is favored, in which 32.4% of the variation in Ka+ can be explained by variation in the recombination rate (Figure 2B; supplementary Table S2). A positive correlation between recombination rate and Ka+ could be due to mutagenic recombination but Wang et al. (2020) found a positive correlation between nucleotide diversity and recombination rate but no correlation between divergence and recombination rate, suggesting that recombination is not mutagenic in Norway spruce. Similarly, genes embedded in gene rich regions are expected to show stronger HRi because more sites are under selection in these genes compared to genes located in gene poor regions, resulting in an expected negative correlation between gene density and the rate of adaptive evolution. We again grouped genes into 10 bins but now based on gene density, with each bin including 5472 scaffolds and 6591 genes on average (supplementary table S4). We observe a negative relationship between Ka+ and gene density with a Spearman’s rank correlation equal to - 0.358 (Figure 2D). Again, a curvilinear model (y =a^bx^) provided a better fit to the data compared to a linear model, showing that gene density explains 10.1% of the variation in Ka+ (Figure 2D; supplementary Table S2). We also calculated correlations between the ratio of non-synonymous to synonymous substitution rates (K_a_/K_s_) and recombination rates or gene density. As expected, K_a_/K_s_ was positively correlated with recombination rate while K_a_/K_s_ was negatively correlated with gene density and for both comparisons a simple linear model was favored (Figure 2A; Figure 2C; supplementary Table S2). All these correlations thus suggest a widespread influence from HRi on the efficiency of adaptive evolution in the coding regions of Norway spruce.

**Figure 2.**
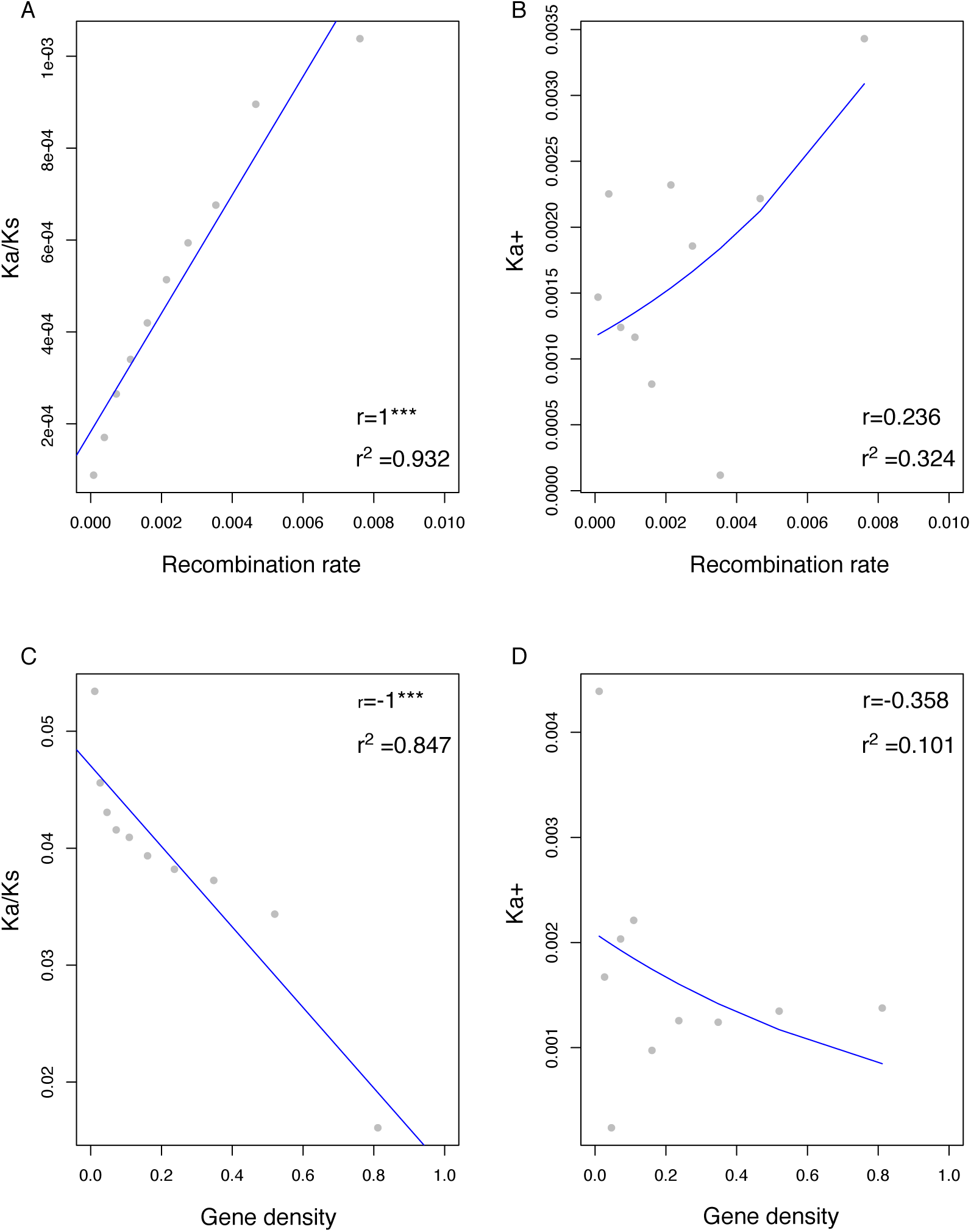
Relations between recombination rate in the x axis and (A) the ratio of non-synonymous to synonymous substitution rates (Ka/Ks) in the y axis, (B) the rate of adaptive amino acid substitutions (Ka+). Relations between gene density and (C) Ka/Ks, (D) Ka+. Each data point has been estimated binning genes. Parameters for each bin can be consulted in the supplementary table S3-S4. The Spearman correlation coefficient (r) and linear/non-linear regression (r^2^) are shown in each plot. Best-fitting regression lines are depicted in blue.

Our results are in accordance with those of Castellano et al. (2016) who found that the rate of adaptive amino acid substitution at a given position of the genome is positively correlated to both the rate of recombination and the mutation rate, and negatively correlated with the gene density of the region in *Drosophila*. Castellano et al (2016) therefore concluded that HRi hampers the rate of adaptive evolution in *Drosophila* and that the variation in recombination, gene density and mutation along the genome affects the impact of HRi. We did not detect any correlations between Ka+ and mutation rate in Norway spruce and thus no support for the hypothesis that genes with high mutation rates adapt at a higher rate than those with low mutation rates. However, this can largely be attributed to the limited outgroup data we have access to which makes it difficult to estimate mutation rates accurately in our study. Moreover, there is evidence that genes with specific functions, such as immune system in *Drosophila* (Obbard et al. 2009), undergo higher rates of adaptive evolution than other genes. The lack of such information in plants suggest that there is a need for more detailed understanding of how gene functions affects the efficiency of adaptive evolution in plants and especially in forest trees.

Variation in effective population size (N_e_) among species also drives variation in the efficacy of selection (Ohta 1973, 1992; Kimura 1983). In species with low N_e_, the impact of positive selection will decrease, leading to lower fixation rates of adaptive mutations and also longer waiting time for beneficial mutations to arise. In contrast, in species with large populations, selection is more effective resulting in higher rates of fixation of adaptive mutations. The end result is that we expect a positive relationship between population size and the rate of adaptive evolution (Gossmann et al. 2010). We calculated N_e_ for the three populations of Norway spruce based on the synonymous site diversity and assuming a mutation rate of 1.1×10^−9^ per site per year and a generation time of 25 years. Compared to the relatively large estimates of N_e_ seen in some flowering plants, e.g. ∼100,000 in *P. tremula* (Ingvarsson 2010), ∼500,000 in *C. grandiflora* (Foxe et al. 2009), ∼832,154 in *H. annuus*, ∼733,133 in *H*. petiolaris (Gossmann et al. 2010), we found a low effective population size in all three populations of Norway spruce (∼46,700 in Finland population, ∼45200 in Sweden-Norway population, and ∼38700 in Central-Europe population), ultimately resulting in the low rates of adaptive evolution we observe in coding regions. Finally, using the folded SFS is expected to yield a greater underestimation of α than using the unfolded SFS (Charlesworth and Eyre-Walker 2008). Using data from a single outgroup individual, as we have done in this study, makes it difficult to infer ancestral states accurately and use of the folded SFS in our calculations can thus be a possible factor contributing to the low estimates for the efficiency of positive selection we observe in Norway spruce.

### Genome-wide scan for regions under positive and balancing selection

Positive selection changes not only the frequency of an advantageous variant but also neighboring polymorphic sites, leaving distinct patterns in the levels of polymorphism and linkage disequilibrium across the genome (Sabeti et al. 2002). Similarly, balancing selection occurs when multiple alleles are maintained at intermediate frequencies in a population, which can result in their preservation over long evolutionary time periods, leading to an excess number of intermediate frequency polymorphisms near a balanced variant (Siewert and Voight 2017). Identifying such genomic signatures through genome-wide scans can help identifying genes or genomic regions that are evolving under the influence of natural selection. This will further help us to better understand the role of natural selection in the evolutionary history of a species and aid in interpreting results for regions previously associated with phenotypes of interest.

We used RAiSD to scan the whole genome for signals of selective sweeps in Norway spruce. The RAiSD algorithm accounts for the expected reduction of variation in the region of a putative selective sweep, the shift in the SFS toward low- and high-frequency derived variants and the emergence of a localized LD pattern characterized by high LD on each side of a beneficial mutation and low LD between loci that are located on different sides of the beneficial allele. We defined the top 0.1% windows of the genome-wide distribution RAiSD values as putative outliers, resulting in a total of 61,756 outlier windows across the whole genome of Norway spruce. These 61,756 windows tag 4208 unique genomic scaffolds and cover a total of 1,594,244 SNPs (Figure 3A; supplementary table S5). We further used a summary statistic-based test, β, to search for signals of long-term balancing selection across the genome for Norway spruce (Siewert & Voight, 2017). We identified a total of 61,451 outlier windows putatively under balancing selection in Norway spruce covering a total of 492,769 variants with those windows (Figure 3B; supplementary table S5).

**Figure 3.**
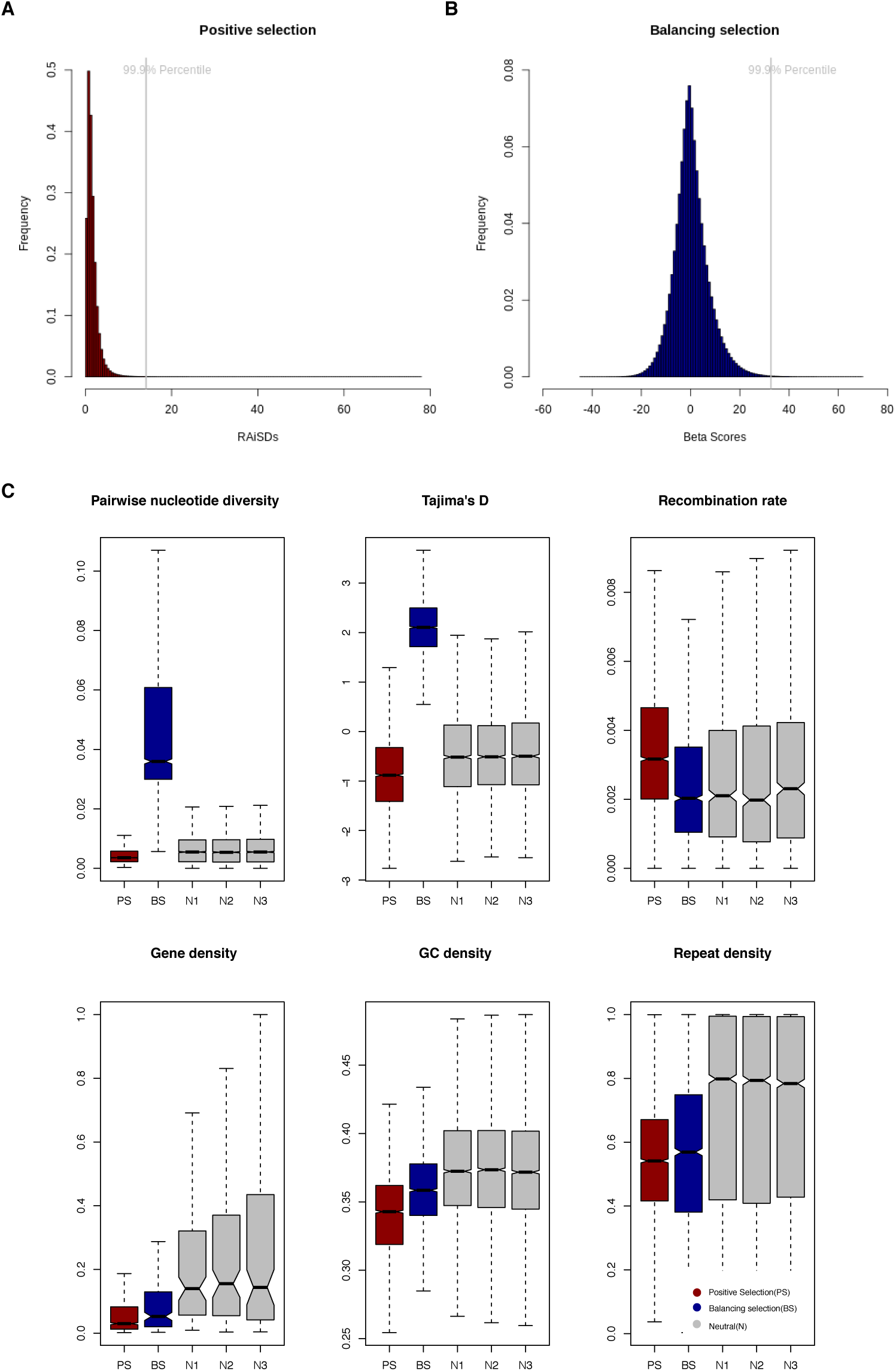
Genome-wide scan to detect outliers under positive selection and balancing selection for Sweden-Norway population (25 individuals). (A) Histogram distribution of RAiSDs to identify outliers under positive selection. (B) Histogram distribution of Beta Scores to identify outliers under balancing selection. (C) Comparisons between outliers identified representing regions under positive (PS) or balancing (BS) selection, with the remaining neutral genomic regions (N1, N2, N3) using a variety of population genetic summary statistics.

In order to understand the impact of positive and balancing selection on genetic variation, we used SNPs within outlier windows that were identified as being under either positive or balancing selection to calculate a variety of population genetic statistics that were then compared to the remaining genomic regions. We also randomly sampled putatively ‘neutral’ scaffolds, creating pseudo-data sets with similar numbers of scaffolds as we observed in the sets of scaffolds under positive and balancing selection to avoid imbalances in the data sets used to calculate population statistics. The fixation of advantageous mutations increase fitness, skew the SFS towards an excess of low and/or high frequency derived alleles and reduce genetic diversity in the vicinity of the selected site (Dutheil 2020), while balancing selection maintains advantageous polymorphisms in populations, leading to an excess number of intermediate frequency polymorphisms and an increased genetic diversity near the balanced variant (De Filippo et al. 2016). In agreement with these predictions, we observe lower pairwise nucleotide diversity (π=0.005) at SNPs under positive selection and higher diversity (π=0.053) at variants under balancing selection compared to the genome-wide average and the randomly sampled “neutral” regions (π=0.006-0.007 (Figure 3C; supplementary Figure S3; supplementary table S6). Similarly, Tajima’s D values are generally positive (2.11) for variants under balancing selection, suggesting an excess of alleles at intermediate frequencies in those regions. As a comparison, both neutral regions and sites under positive selection showed negative Tajima’s D values, where the negative values (−0.423 to -0.445) of neutral regions are indicative of a demographic history of Norway spruce with population expansion after recent bottleneck (Wang et al. 2020), while the more negative value (−0.849) for sites under positive selection suggested an even greater abundance of rare alleles compared to that in neutral regions. We found that regions under positive selection had higher recombination rates and lower gene densities compared to neutral regions. These results are in accordance with our previous results that we confirmed a positive correlation between the adaptive rate (Ka+) and recombination rate and the negative correlation between adaptive rate and gene density, both of which are mediated by Hill-Robertson interference (Figure 2). However, compared with neutral regions, we observed lower recombination rates in regions under balancing selection, possibly because the identification of a locus with an old balanced polymorphism is facilitated by low recombination rates that results in the genealogical histories of adjacent SNPs to be more strongly correlated. In these cases, the ability to pinpoint the actual target of selection will also be reduced because larger segments of the genome will be affected (Tian et al. 2002).Moreover, we found that both GC density and repeat density were lower in regions under positive and balancing selection compared to neutral regions. Apuli et al. (2020) found a significantly positive correlation between gene density and GC contents, and a significantly negative correlation between repeat density and LD-based recombination rates in European aspen (*Populus tremula*). A similar pattern in Norway spruce would explain the low GC content and repeat density for sites under positive selection and the low GC content for sites under balancing selection that we observed in Norway spruce. However, although we detect low repeat densities around sites under balancing selection, there is evidence of balancing selection on transposable elements (TEs). For instance, Chen et al. (2017) showed that the heat-shock protein *Hsp*90 is found only in a heterozygote state and seems to display latitudinal variation (Bourgeois and Boissinot 2019). The role of repeats in regions undergoing balancing selection in Norway spruce is thus worth investigating in greater detail in future studies.

### Genes under positive and balancing selection

To assess whether there were any specific biological functions that were significantly overrepresented among the genes located in regions identified as undergoing either positive (197 genes) or long-term balancing selection (13 genes), we performed gene ontology (GO) enrichment analysis. We identified 27 significantly enriched GO categories for genes under positive selection, which all belong to different molecular functions (supplementary table S7). These GO clusters were primarily associated with transporter activity (transmembrane, substrate-specific), binding process (heterocyclic and organic cyclic compound, small molecule, nucleotide, nucleoside phosphate, cofactor, isoprenoid, monocarboxylic acid, identical protein, hormone and coenzyme), ATPase activity, hydrolase activity, protein homodimerization activity, catalytic activity and nucleoside-triphosphatase activity. We only detected one significant GO terms for the candidate genes under long-term balancing selection with FDR corrected P<0.1, associated with molecular function of protochlorophyllide reductase activity. In addition, other GO categories for candidate genes under long-term balancing selection that were significant before false-discovery correction include both molecular function of glucan exo-1,3-beta-glucosidase activity, oxidoreductase activity transferase activity, dCMP deaminase activity, and biological process involved in skeletal muscle contraction and negative regulation of humoral immune response (supplementary table S7).

**Figure 4.**
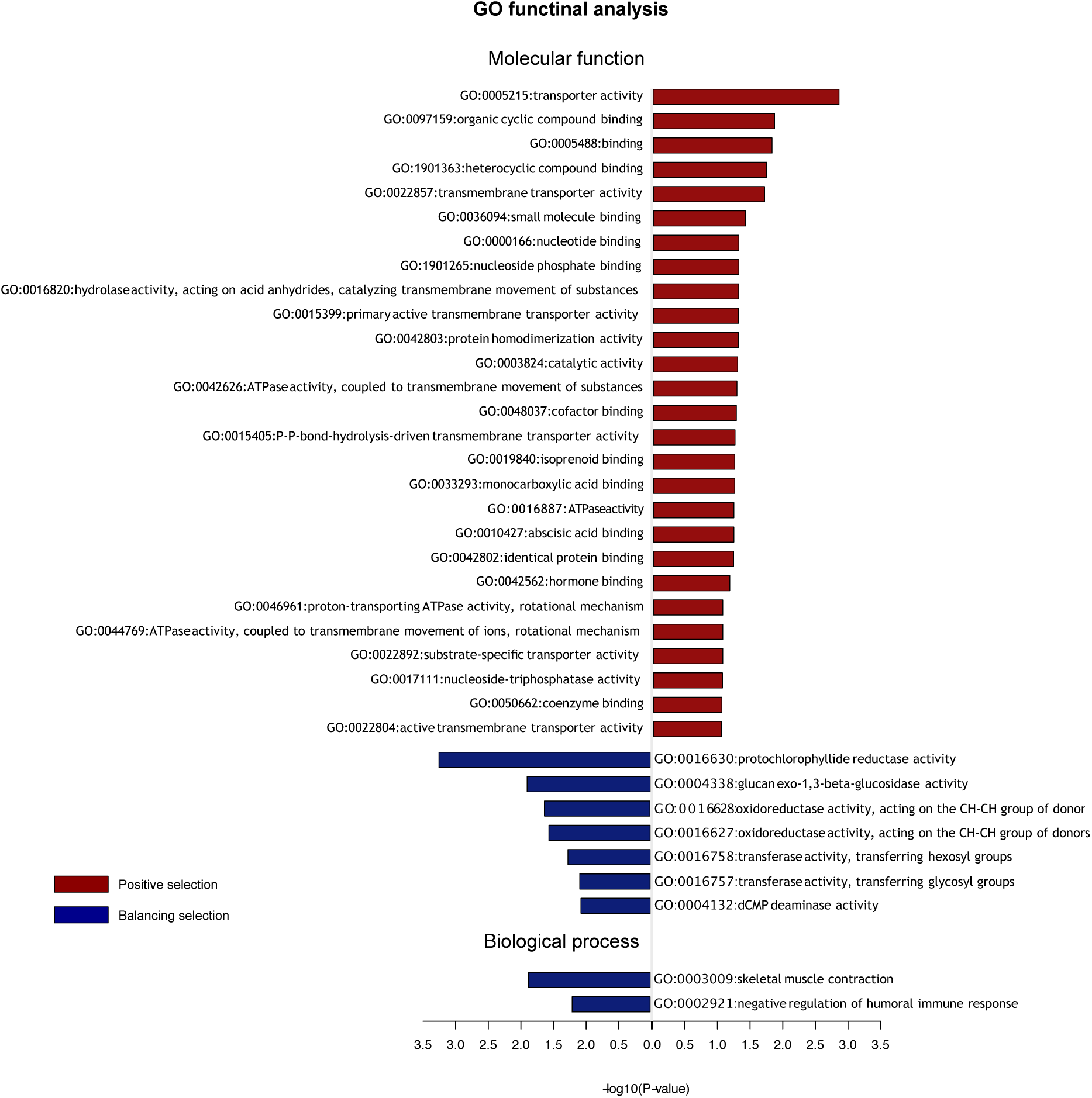
Enriched GO categories for genes under positive selection and balancing selection.

## Conclusion

In this population genomic survey, we use whole genome re-sequencing data from 34 individuals of Norway spruce, spanning most of the natural distribution of the species. We first evaluate the efficacy of both purifying and positive selection across the whole genome of *P. abies*. Our results show that negative selection is limited to coding regions, with both introns and intergenic regions approaching 100% of mutations falling in the N_e_s range nearly neutral mutations. This is consistent with earlier research in Norway spruce. For instance, Chen et al. (2016) found non-synonymous SNPs tended to have both lower frequency differences and lower F_st_ values than silent ones between the Alpine and Fennoscandian domains, suggesting the presence of purifying selection in Norway spruce. Further, Wang et al. (2020) indicated that background selection may be limited in *P. abies*, by showing a very weak negative correlation between nucleotide diversity and gene density. As a contrast, positive selection seems very strong in non-coding regions in the Norway spruce genome, suggesting the great importance of regulatory changes in evolution (King and Wilson 1975; Carroll et al. 2001). We did find rare evidence of positive selection in coding regions in Norway spruce, which mirrors results on patterns of protein adaptive in 11 species of soft pines (Eckert et al. 2013). However, our conifer results are in stark contrast to patterns observed in many outcrossing flowering plants, e.g. *P. tremula* (Ingvarsson 2010), *C. grandiflora* (Slotte et al. 2010), and sunflowers (Strasburg et al. 2009). We further analyzed which factors are important for determining the efficacy of protein adaptation in Norway spruce. We observe a positive correlation between the rate of adaptive evolution and recombination rates and a negative correlation between the rate of adaptive evolution and gene density which both suggest a widespread influence from Hill-Robertson interference. Castellano et al. (2016) also found such correlations and concluded that HRi hampers the rate of adaptive evolution in *Drosophila*. We finally scanned the Norway spruce genome to identify potential outlier regions evolving under either positive and balancing selection, and compared population statistics for those outliers with neutral regions. Gene ontology enrichment analysis for genes located in regions identified as undergoing either positive or long-term balancing selection also highlighted specific molecular functions and biological processes in that appear to be targets of selection in Norway spruce. This study constitutes one of the first to find convincing evidence of natural selection within both coding and non-coding genome for a conifer species. Future studies should aim to use even more broadly sampled populations of the species to further extend the analyses on the importance of adaptive evolution throughout the evolutionary history of Norway spruce.

## Acknowledgements

The research has been funded by grants from the Knut and Alice Wallenberg Foundation (Norway spruce genome project) and the Swedish Foundation for Strategic Research (SSF, Grant No. RBP14-0040). Data generation was supported by Science for Life Laboratory and the National Genomics Infrastructure (NGI) which provided access to massive parallel sequencing. All analyses were performed on resources pro-vided by the Swedish National Infrastructure for Computing (SNIC) at Uppsala Multidisciplinary Centre for Advanced Computational Science (UPPMAX) under the projects b2012141, SNIC 2017/1-438, SNIC 2018/3-529 and uppstore2017066. X.W. was supported by a scholarship from the Chinese Scholarship Council (CSC).

