## Supplementary Figures for "Quantifying adaptive evolution and the effects of natural selection across the Norway spruce genome"

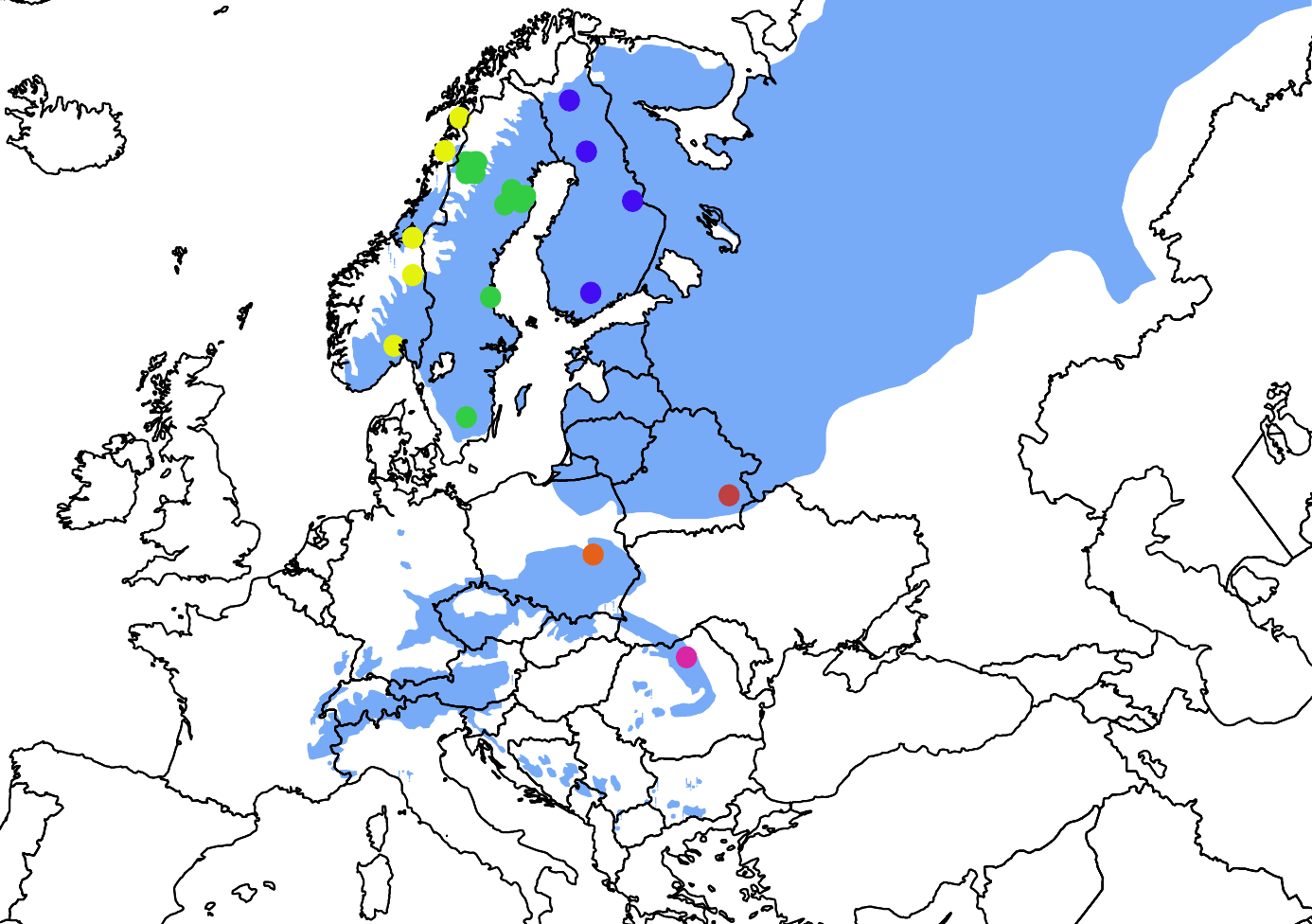

**Figure S1**. Geographic distribution of whole-genome re-sequenced 34 individuals. Individuals of *P.abies* from Norway (yellow), Sweden (green), Finland (blue), Poland (orange), Belarus (red), and Romania (pink) are shown in circles with different colors.

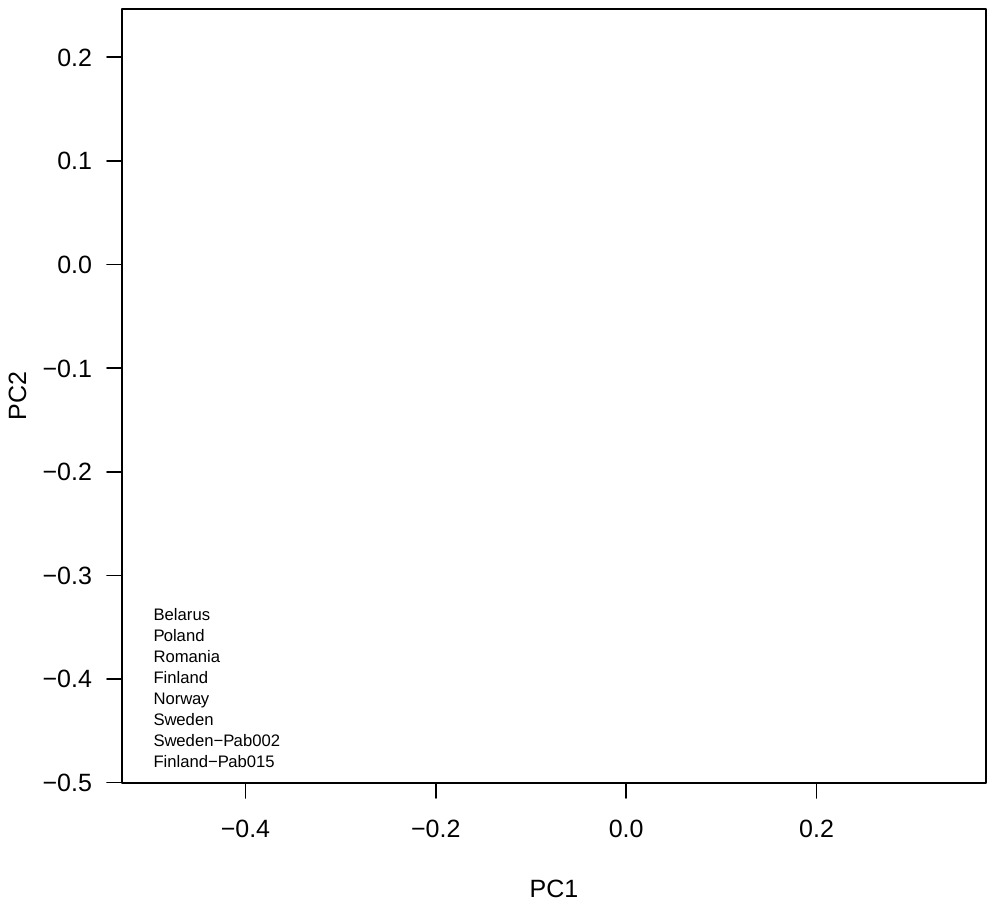
**Figure S2**. Population structure of *P. abies*. PCA plot calculated from 4fold synonymously SNPs among 33 individuals by PCAngsd. Individual Pab034 was removed because it was found to be highly related with individual Pab033 (Wang et al. 2020).

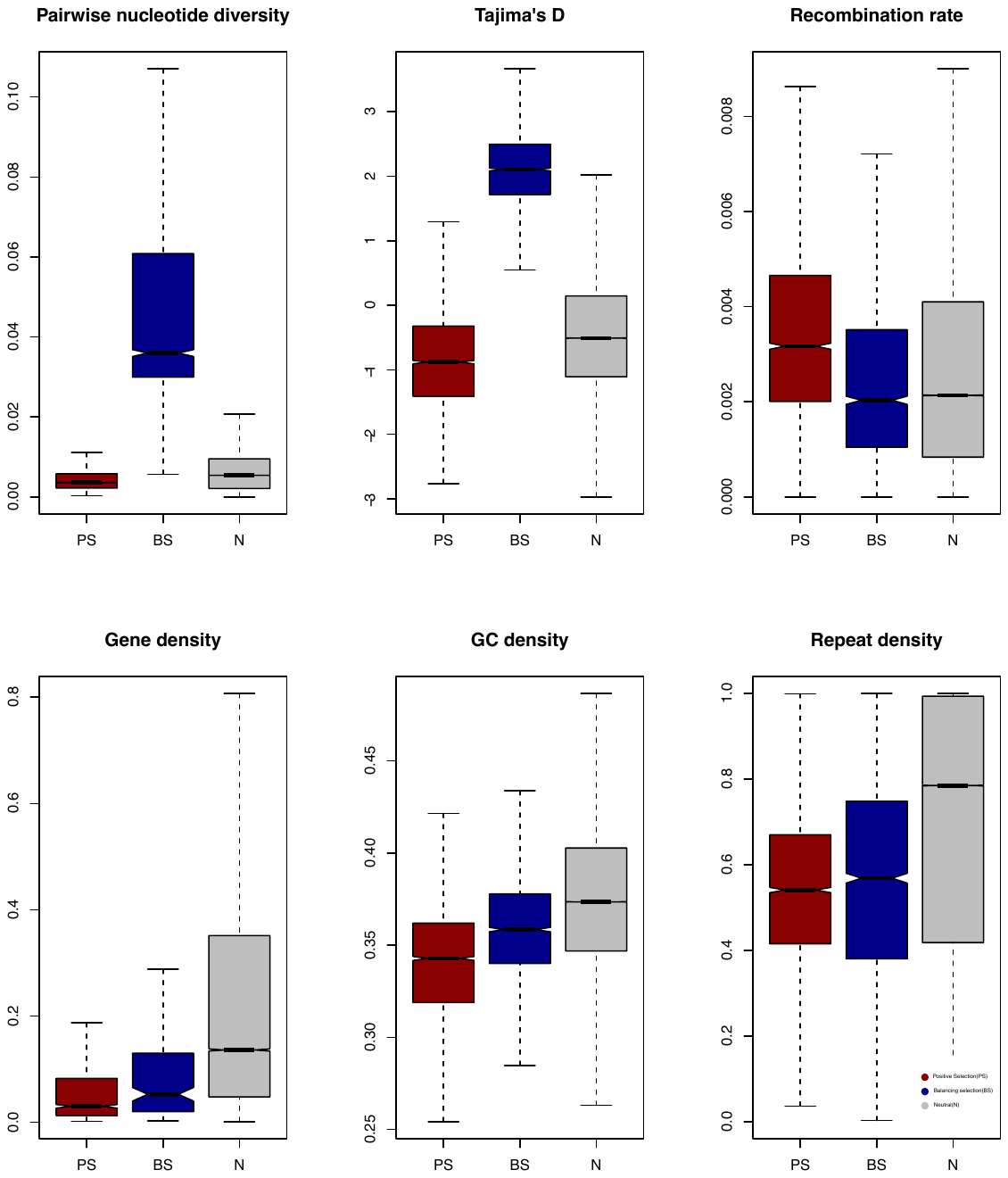
**Figure S3.** Comparisons between outliers identified representing regions under positive (PS) or balancing (BS) selection, with the remaining neutral genomic regions (N) using population genetic summary statistics for Sweden-Norway population(25inds).
